# TAK1 regulates skeletal muscle mass, hypertrophic signaling, and metabolic homeostasis in male and female mice

**DOI:** 10.64898/2026.02.26.708345

**Authors:** Meiricris Tomaz da Silva, Aniket S. Joshi, Anirban Roy, Troy A. Hornberger, Ashok Kumar

## Abstract

Skeletal muscle is the most abundant tissue in the human body and is essential for locomotion and the regulation of whole-body metabolism. The maintenance of skeletal muscle mass is essential for health, yet the molecular and signaling mechanisms that control skeletal muscle mass remain poorly understood. Transforming growth factor-β-activated kinase 1 (TAK1) is a key signaling protein that regulates multiple intracellular pathways. Recent studies have demonstrated that TAK1 is a critical regulator of skeletal muscle mass. However, the mechanisms by which TAK1 regulates muscle mass and whether its role is sex dependent remain incompletely understood. In this study, we show that targeted inactivation of TAK1 induces muscle atrophy more rapidly in male than in female mice. Loss of TAK1 activity also abolished mechanical overload-induced phosphorylation of p70S6K and rpS6, and the induction of myofiber hypertrophy in both sexes. RNA-Seq analysis further revealed that TAK1 inactivation in skeletal muscle disrupts the gene expression of various molecules involved in catabolic processes, calcium signaling, muscle structure development, and aerobic respiration. Moreover, TAK1 inactivation impairs fatty acid oxidation and promotes lipid accumulation in skeletal muscle of adult mice in a sex-independent manner. Collectively, our findings demonstrate that TAK1 regulates skeletal muscle mass and growth by coordinating distinct intracellular pathways in both male and female mice.

## Introduction

Skeletal muscle is one of the largest organs in the body and performs several critical functions, including heat generation, respiration, voluntary movement, maintenance of posture, and regulation of whole-body metabolism. Skeletal muscle also exhibits remarkable plasticity, enabling dynamic structural and metabolic adaptations in response to environmental and physiological cues. For example, resistance exercise and adequate nutritional intake promote an increase in myofiber size, resulting in muscle hypertrophy. In contrast, prolonged inactivity and various chronic disease conditions reduce myofiber diameter, leading to skeletal muscle atrophy. Muscle atrophy is a serious complication that contributes substantially to morbidity and mortality (1, 2), highlighting the importance of understanding the cellular and molecular mechanisms that regulate skeletal muscle mass to preserve overall health and improve quality of life.

Skeletal muscle mass is regulated by the coordinated activation of multiple signaling pathways. Among these, the IGF1-Akt-mTORC1 axis is one of the best-characterized pathways promoting myofiber hypertrophy by increasing the rate of protein synthesis (3-5). Activated mTORC1 phosphorylates eIF4E-binding protein 1 (4E-BP1), causing its dissociation from the translation initiation factor eIF4E. This release enables eIF4E to associate with eIF4G, facilitating formation of the eIF4F complex and cap-dependent initiation of translation (4, 6). In addition, mTORC1 phosphorylates p70 ribosomal protein S6 kinase beta-1 (p70S6K1), which enhances protein synthesis by phosphorylating ribosomal protein S6 (rpS6), a component of the 40S ribosomal subunit (4, 7-11). Growth factors and resistance exercise activate rapamycin-sensitive mTORC1 signaling, leading to translation initiation and a net increase in protein synthesis in skeletal muscle (3, 12-15).

Intriguingly, accumulating evidence suggests that the prolonged elevation in muscle protein synthesis following a bout of resistance exercise is largely rapamycin-insensitive (16, 17). Moreover, inhibition of mTORC1 through inducible deletion of Raptor in skeletal muscle does not block the increase in protein synthesis induced by passive stretch or chronic mechanical overload, suggesting that mTORC1-independent mechanisms contribute to mechanically stimulated protein synthesis (18-20). Recent studies have also shown that constitutive activation of mTORC1 can lead to skeletal muscle atrophy and myopathy without significantly enhancing protein synthesis (21-24), indicating that mTORC1-independent pathways play important roles in regulating skeletal muscle mass and growth under various conditions.

Transforming growth factor-β-activated kinase 1 (TAK1) is a key signaling molecule that mediates activation of multiple intracellular pathways in response to growth factors and cytokines (23, 25, 26). Our recent studies have identified TAK1 as a major regulator of skeletal muscle mass. We demonstrated that germline, muscle-specific inactivation of TAK1 in mice results in perinatal lethality (27). Importantly, tamoxifen-inducible deletion of TAK1 in adult mice leads to muscle atrophy, mitochondrial dysfunction, oxidative stress, and degeneration of neuromuscular junctions (NMJs), phenotypes reminiscent of aged skeletal muscle (27-29). We further showed that activation of TAK1 enhances protein synthesis and promotes muscle growth in adult mice (28). However, whether TAK1 regulates skeletal muscle mass in a sex-dependent manner remains unknown. In addition, the signaling and molecular mechanisms by which TAK1 promotes myofiber growth in response to mechanical overload have not been elucidated.

In this study, we used tamoxifen-inducible TAK1 knockout mice to examine the role of TAK1 in the regulation of skeletal muscle mass in adult male and female mice and to define the mechanisms through which TAK1 enhances myofiber growth in response to mechanical overload. Our results show that while the onset is delayed in females, TAK1 inactivation ultimately causes muscle atrophy in both sexes. We further demonstrate that the loss of TAK1 activity abolished mechanical overload-induced phosphorylation of p70S6K and rpS6, and the induction of myofiber hypertrophy in both sexes. RNA-seq analysis revealed that TAK1 inactivation disrupts calcium signaling, endoplasmic reticulum (ER) stress-induced unfolded protein response (UPR), aerobic respiration, and fatty acid metabolism in skeletal muscle of adult mice.

## Results

### Targeted inactivation of TAK1 reduces muscle mass selectively in male mice

We have previously reported that targeted inactivation of TAK1 in adult mice causes loss of body weight and skeletal muscle mass (27, 29). However, it was not clear whether TAK1 affects these parameters in a sex-dependent manner. By crossing Tak1^fl/fl^ and HSA-MCM mice, we generated tamoxifen-inducible muscle-specific Tak1-knockout (Tak1^mKO^) mice and littermate control mice (Tak1^fl/fl^) mice as described (27). Next, 10-week-old male and female Tak1^fl/fl^ and Tak1^mKO^ mice were given intraperitoneal injection of tamoxifen for four consecutive days. After a washout period of two days, the mice were fed a tamoxifen containing chow for the entire duration of the experiment. The mice were then analyzed for body weight. There was a significant reduction in the body weight of male Tak1^mKO^ mice compared to male Tak1^fl/fl^ mice within 15 days which persisted till day 30 of the start of tamoxifen injection (**Fig. 1A**). By contrast, there was no significant difference in the body weight of female Tak1^fl/fl^ and Tak1^mKO^ mice on day 15 after tamoxifen administration. While there was a trend towards a decrease in body weight of female Tak1^mKO^ compared with female Tak1^fl/fl^ mice on day 30 after the start of tamoxifen treatment, this difference was not statistically significant (**Fig. 1B**). Next, the mice were euthanized, and the wet weight of individual hind limb muscle was measured. Results showed that there was a significant reduction in the absolute wet weight of tibialis anterior (TA), soleus (SOL), and gastrocnemius (GA) muscle of male Tak1^mKO^ mice compared with male Tak1^fl/fl^ mice (**Fig. 1C**). In contrast, no significant differences in the absolute wet weight of TA, SOL, or GA muscles were observed between female Tak1^fl/fl^ and Tak1^mKO^ mice after 30 days of inactivation of TAK1 (**Fig. 1C**). Furthermore, there was a significant reduction in the wet weight of TA and GA muscle, but not soleus muscle, normalized by body weight of male Tak1^mKO^ mice compared with male Tak1^fl/fl^ mice, whereas normalized muscle weights were comparable between female Tak1^fl/fl^ and Tak1^mKO^ mice for all muscles analyzed (**Fig 1D**).

**FIGURE 1.**
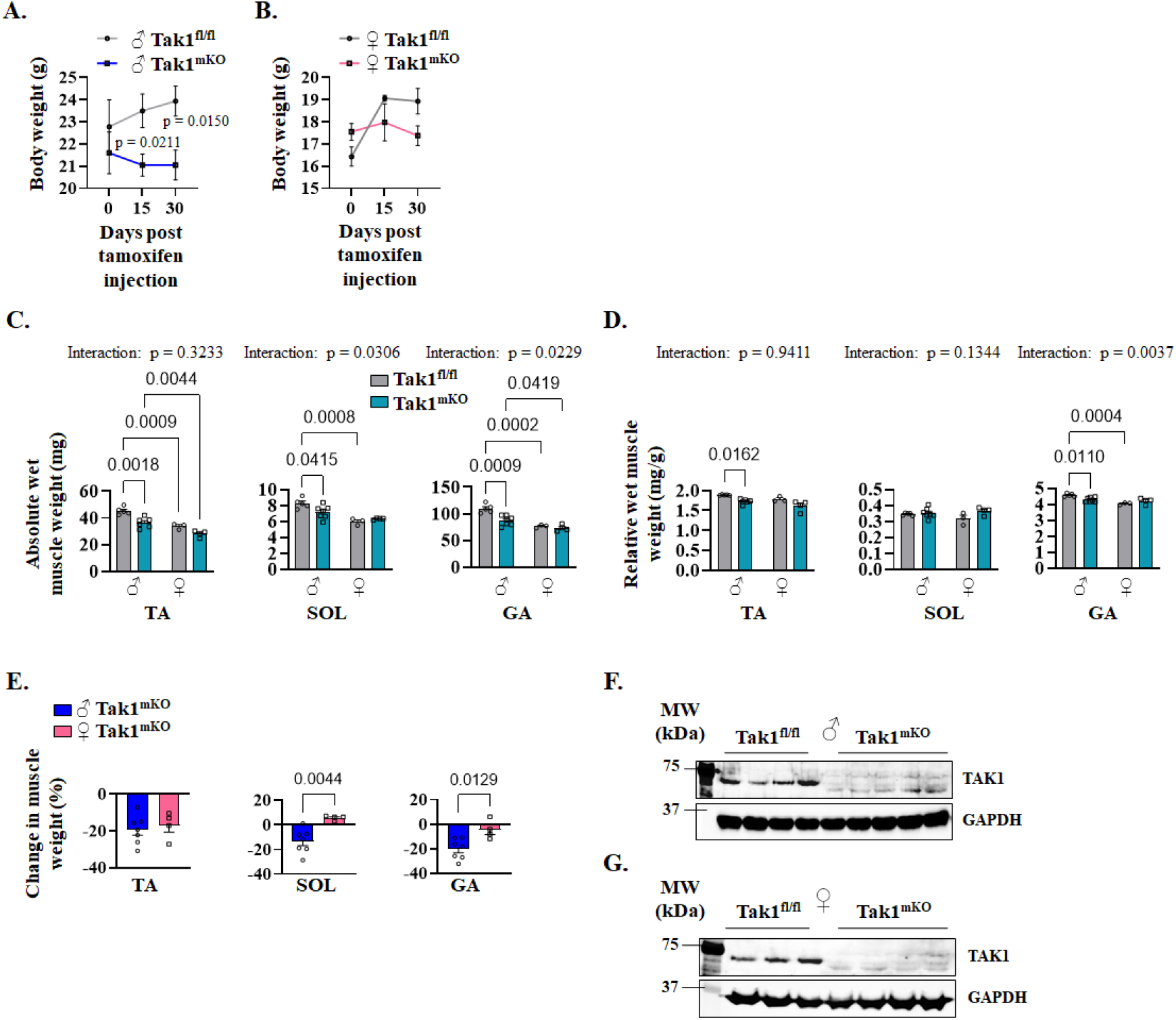
Effect of inactivation of TAK1 on skeletal muscle mass in male and female mice. **(A)**Body weight (BW) of Tak1^fl/fl^ and Tak1^mKO^ male mice on day 0, 15 and 30 after tamoxifen injection. **(B)** BW of Tak1^fl/fl^ and Tak1^mKO^ female mice on day 0, 15 and 30 after tamoxifen injection. **(C)** Absolute wet weights, and **(D)** wet weights of Tibialis anterior (TA), soleus (SOL), and gastrocnemius (GA) muscle relative to BW of adult male and female Tak1^fl/fl^ and Tak1^mKO^ mice on day 30 after tamoxifen injection. **(E)** Change in wet weight of TA, SOL and GA muscle of male or female Tak1^mKO^ mice compared to corresponding sex Tak1^fl/fl^ mice. . Immunoblots showing levels of TAK1 protein in GA muscle of **(I)** male and **(F)** female Tak1^fl/fl^ and Tak1^mKO^ mice. n=3-7 mice per group. All data are presented as mean ± SEM. The data were analyzed with an unpaired t-test (A, B, E), two-way ANOVA and Tukey’s multiple comparison test (C, D), and the resulting p-values for the interactions, and pairwise comparisons are displayed in the graphs.

Further analysis showed that the change in wet muscle weight in response to TAK1 inactivation was similar in the TA muscle of both males and females. In contrast, the percentage loss of wet weight of soleus and GA muscle was significantly less in female Tak1^mKO^ mice compared to male Tak1^mKO^ mice (**Fig. 1E**). Because TAK1 inactivation did not alter muscle mass in female mice, the reduction observed in males apopears to reflects a sex-specific effect rather than a general response to TAK1 deletion. By performing western blot, we confirmed that treatment with tamoxifen considerably reduced the levels of TAK1 in the GA muscle of both male and female Tak1^mKO^ mice compared to corresponding Tak1^fl/fl^ mice (**Fig. 1F, G**).

### Targeted inactivation of TAK1 reduces myofiber size in adult mice

We next generated transverse sections of TA and soleus muscle of male and female Tak1^fl/fl^ and Tak1^mKO^ mice followed by staining with Hematoxylin and Eosin (H&E) or anti-dystrophin (to mark myofiber boundary) and DAPI (**Fig. 2A, 2B**). Analysis of the H&E staining section showed that there was no overt effect of TAK1 inactivation on TA or soleus structure except that myofiber cross-sectional area was reduced (**Fig. S1**). Quantitative analysis showed that there was significant reduction in the average myofiber CSA in TA muscle of both male and female Tak1^mKO^ mice compared with their respective Tak1^fl/fl^ mice (**Fig. 2C**) with a comparable percentage decrease between sexes (**Fig. 2D**). Our morphometric analysis also showed that there was a significant reduction in the average myofiber CSA in the soleus muscle of both male and female Tak1^mKO^ compared to corresponding Tak1^fl/fl^ mice (**Fig. 2E**). However, the percentage reduction in myofiber CSA was significantly smaller in female Tak1^mKO^ than in male Tak1^mKO^ mice (**Fig. 2F**). Our analysis showed that the proportion of smaller myofibers was markedly increased in the TA and soleus muscles of male and female TAK1^mKO^ mice compared with corresponding male or female Tak1^fl/fl^ mice (**Fig. 2G, H**). Collectively, these results indicate that TAK1 deletion leads to muscle atrophy in both male and female mice, with a more pronounced in males, particularly in the soleus muscle.

**FIGURE 2.**
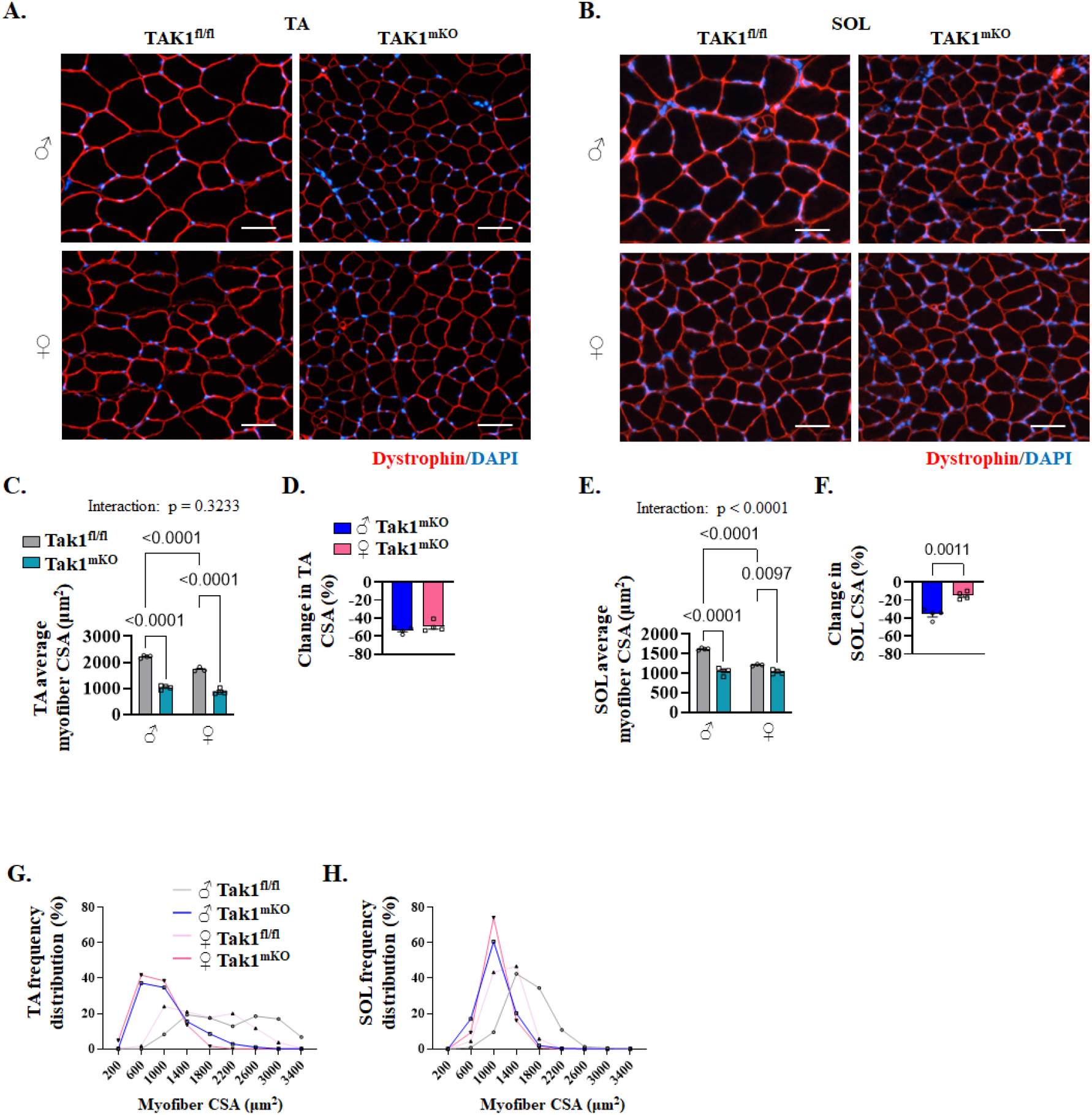
Effect of targeted inactivation of TAK1 on myofiber size in male and female mice. Representative photomicrographs of **(A)** TA, and **(B)** soleus (SOL) muscle sections from male and female Tak1^fl/fl^ and Tak1^mKO^ mice on day 30 after tamoxifen injection, after performing anti-dystrophin and DAPI staining, Scale bar, 50 μm. **(C)** Quantification of average myofiber cross-sectional area (CSA) in TA muscle of male and female Tak1^fl/fl^ and Tak1^mKO^ mice. **(D)** Percentage change in average myofiber CSA in TA muscle of male and female Tak1^mKO^ mice compared to sex-matched Tak1^fl/fl^ mice. **(E)** Quantification of average myofiber CSA in soleus muscle of male and female Tak1^fl/fl^ and Tak1^mKO^ mice. (**F)** Percentage change in average myofiber CSA in soleus muscle of male and female Tak1^mKO^ mice compared to sex-matched Tak1^fl/fl^ mice. **(G)** Histograms showing frequency distribution of myofiber CSA in TA muscle of male and female Tak1^fl/fl^ and Tak1^mKO^ mice. **(H)** Histograms showing frequency distribution of myofiber CSA in soleus muscle of male and female Tak1^fl/fl^ and Tak1^mKO^ mice. n=3-4 mice per group. All data are presented as mean ± SEM. The data were analyzed with two-way ANOVA and Tukey’s multiple comparison test (C, D), unpaired t-test (E, F), and the resulting p-values for the interactions, and pairwise comparisons are displayed in the graphs.

### Inactivation of TAK1 inhibits mechanical overload-induced muscle growth in adult male and female mice

We first investigated how the levels of TAK1 are affected in response to mechanical overload (MOV). About 10-12-week-old wild-type mice were subjected to sham or bilateral MOV surgery and the plantaris muscle was isolated on day 7 followed by performing western blot. Results showed that the levels of both phosphorylated and total TAK1 protein were significantly increased in overloaded plantaris muscle of both male and female mice compared to corresponding controls (**Fig. 3A-D**). To understand the role of TAK1 in MOV-induced muscle growth, 10-week-old male and female Tak1^fl/fl^ and Tak1^mKO^ were treated with tamoxifen for 4 consecutive days. After a 2-day washout period, mice underwent sham or bilateral MOV surgery. The mice were euthanized on day 14 following MOV surgery and the plantaris muscles were isolated and analyzed by morphometric methods. There was a significant increase in the plantaris muscle wet weight normalized by body weight in male Tak1^fl/fl^ and Tak1^mKO^ mice subjected to MOV surgery compared to those subjected to sham surgery. However, the normalized wet weight of plantaris muscle was significantly less in male Tak1^mKO^ mice compared with male Tak1^fl/fl^ mice following MOV (**Fig. 3E**). Interestingly, MOV led to a significant increase in the normalized wet weight of plantaris muscle of female Tak1^fl/fl^, but not of female Tak1^mKO^ mice (**Fig. 3F**). We next generated transverse sections of plantaris muscle and performed H&E or anti-dystrophin staining followed by measuring average myofiber CSA. There was no overt structural difference in the plantaris muscle of Tak1^fl/fl^ and Tak1^mKO^ mice subjected to sham or MOV surgery (**Fig. S2**). Morphometric analysis of anti-dystrophin-stained sections showed that MOV led to a significant increase in average myofiber CSA in both male and female Tak1^fl/fl^ mice. The average myofiber CSA of the plantaris muscle was smaller in sham-operated male and female Tak1^mKO^ mice compared to corresponding Tak1^fl/fl^ mice. Importantly, there was no significant increase in average myofiber CSA in male or female Tak1^mKO^ in response to MOV (**Fig. 3G-J**). Distribution of myofiber CSA also showed that the MOV led to an increase in the proportion of myofiber with higher CSA in the plantaris muscle of male and female Tak1^fl/fl^ mice, but not Tak1^mKO^ (**Fig. 3K, L**). These results suggest that TAK1 is essential for overload-induced myofiber growth in both male and female mice.

**FIGURE 3.**
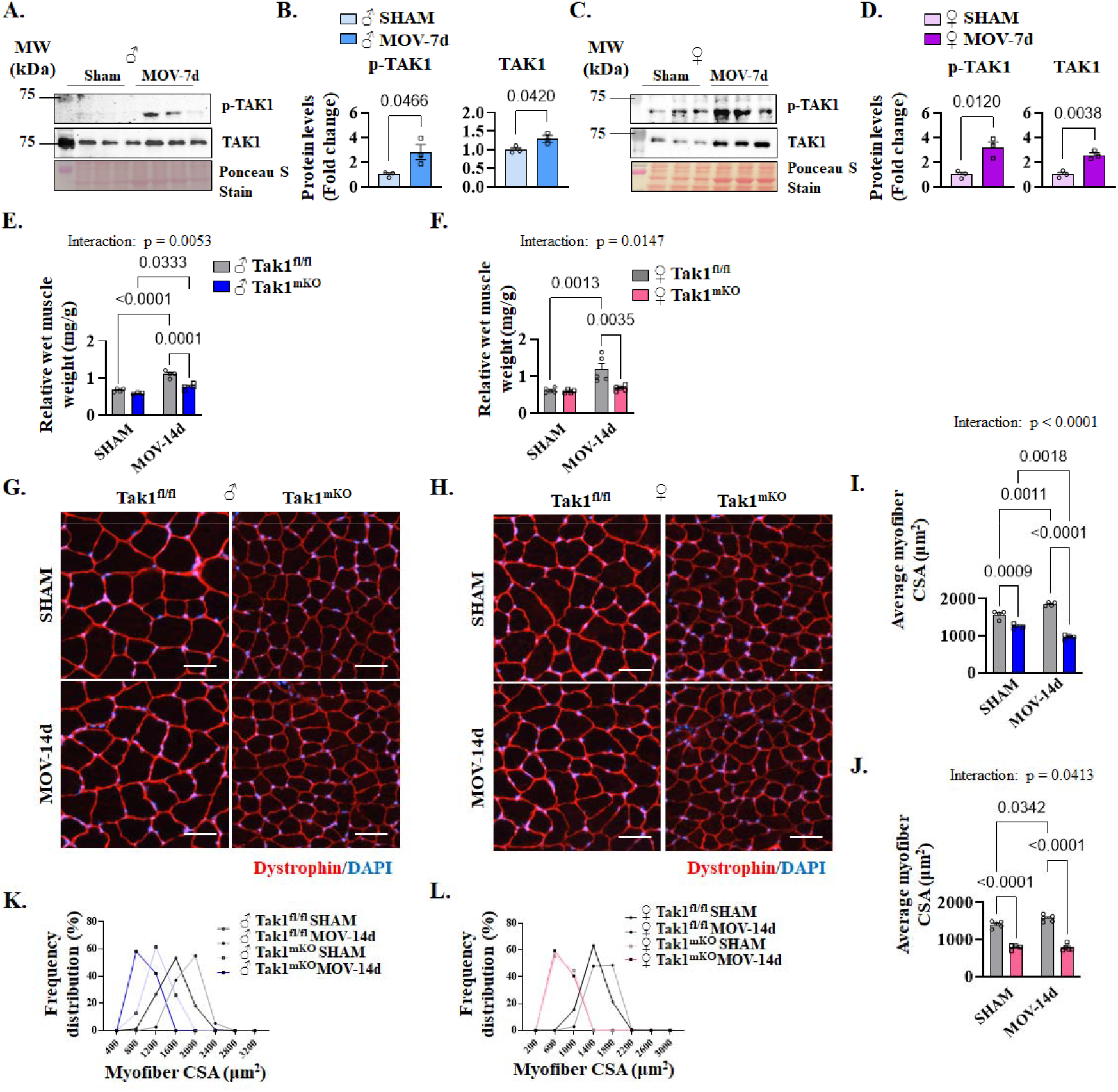
Effect of targeted inactivation of TAK1 on mechanical overload-induced muscle hypertrophy in adult mice. **(A)**Representative immunoblots, and **(B)** densitometry analysis of levels of phosphorylated and total TAK1 protein in plantaris muscle of 12-week-old male wild-type mice on day 7 after performing sham or bilateral mechanical overload (MOV) surgery. **(C)** Representative immunoblots, and **(D)** densitometry analysis of levels of phosphorylated and total TAK1 protein in plantaris muscle of 12-week-old female wild-type mice on day 7 after performing sham or bilateral MOV surgery. Quantification of relative wet weight of plantaris muscle of **(E)** male and **(F)** female Tak1^fl/fl^ and Tak1^mKO^ mice on day 14 after performing sham or MOV surgery. Representative anti-dystrophin and DAPI-stained photomicrographs of plantaris muscle sections of **(G)** male, and **(H)** female Tak1^fl/fl^ and Tak1^mKO^ mice after 14 d of performing sham or bilateral MOV surgery. Scale bar, 50 μm. **(I)** Quantification of average myofiber cross-sectional area (CSA) in control (sham surgery) and 14d-overloaded (MOV surgery) plantaris muscle of **(I)** male and **(J)** female Tak1^fl/fl^ and Tak1^mKO^ mice. Histograms showing frequency distribution of myofiber CSA in control and 14d-overloaded plantaris muscle of **(K)** male, and **(L)** female Tak1^fl/fl^ and Tak1^mKO^ mice. n=3-5 mice per group. All data are presented as mean ± SEM. The data were analyzed with an unpaired t-test (B, D), two-way ANOVA and Tukey’s multiple comparison test (E, F, I, J), and the resulting p-values for the interactions, and pairwise comparisons are displayed in the graphs.

### TAK1 regulates the mechanical overload-induced phosphorylation of p70S6K and rpS6

To understand the signaling mechanisms by which TAK1 augments myofiber growth in response to mechanical overload, we measured the levels of phosphorylated and total Akt, mTOR, p70S6K, 4E-BP1, and Rsp6 proteins in plantaris muscle following 7 days of MOV in both sexes. MOV led to a significant increase in the levels of phosphorylated Akt in Tak1^fl/fl^ mice compared with sham controls. Total Akt protein levels were also increased following MOV in both Tak1^fl/fl^ and Tak1^mKO^ mice. Notably, after MOV, total Akt levels were significantly higher in Tak1^mKO^ mice compared to Tak1^fl/fl^ mice (**Fig. 4A-C**). Similarly, MOV resulted in a significant increase in the levels of phosphorylated mTOR and 4E-BP1 in both Tak1^fl/fl^ and Tak1^mKO^ mice compared with their respective sham controls. Total mTOR and 4E-BP1 protein levels also showed a comparable pattern, with increased levels observed in both genotypes following MOV.

**FIGURE 4.**
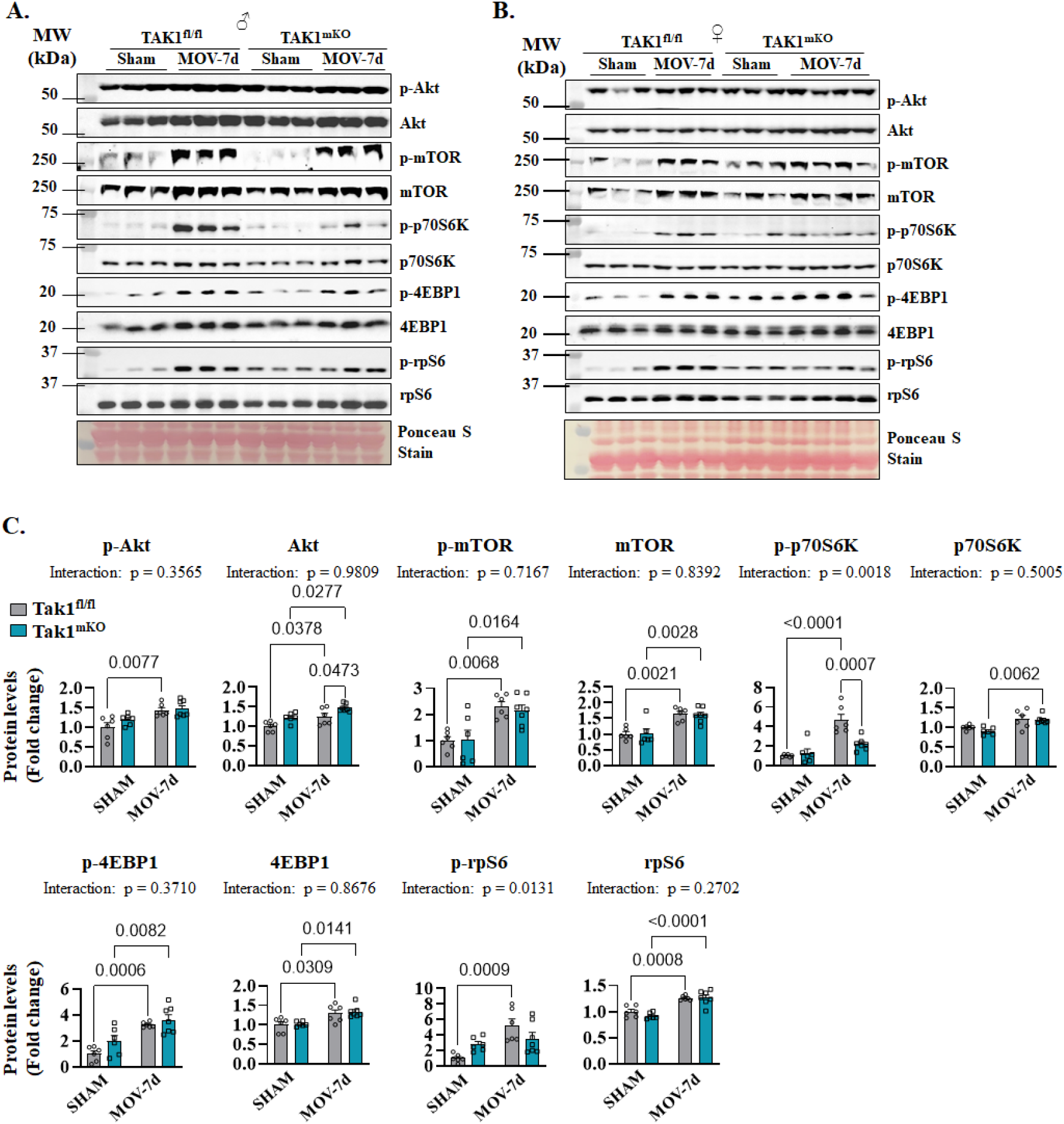
Effect of targeted inactivation of TAK1 on the phosphorylation of the components of Akt-mTOR signaling in male mice. **(A), (B)** Immunoblots and **(C)** densitometry analysis of phosphorylated and total levels of Akt, mTOR, p70S6K, 4EBP1, and rpS6 protein in plantaris muscle of male and female Tak1^fl/fl^ and Tak1^mKO^ mice on day 7 after performing sham or MOV surgery. n=6-7 mice per group. All data are presented as mean ± SEM. The data were analyzed with two-way ANOVA and Tukey’s multiple comparison test, and the resulting p-values for the interactions, and pairwise comparisons are displayed in the graphs.

The above resuls indicated that Tak1 was not necessary for the MOV-induced changes in total and phosphorylated forms of Akt, mTOR and 4E-BP1. In contrast, Tak1 appears to be essential for the phosphorylation of p70S6K and rpS6. Specifically, in Tak1^fl/fl^ mice, MOV elicited a robust increase in phosphorylated p70S6K and rpS6, whereas this response was largely abolished in the absence of Tak1 (**Fig. 4A-C**). Importantly, the differential responses observed between Tak1^fl/fl^ and Tak1^mKO^ mice could not be explained by differences in the response at the total protein level of p70S6K and rpS6. Hence, it appears that Tak1 plays a critical role in the signaling pathway through which MOV stimulates p70S6K and rpS6 phosphorylation.

We also performed quantification of western blot data separately for both sexes. Again, we found similar changes in all these proteins except that the basal level of 4-EBP1 protein was significantly higher in female Tak1mKO mice compared to correspoing female Tak1fl/fl mice (**Fig. S3**). Collectively, these results suggest that TAK1 is required for overload-induced activation of p70S6K and rpS6, while activation of Akt, mTOR, and 4E-BP1 occurs independently of TAK1.

### Targeted inactivation of TAK1 disrupts the gene expression of molecules involved in calcium signaling and ER stress-induced UPR

To further understand the mechanisms by which TAK1 regulates skeletal muscle mass, we performed bulk RNA-seq analysis of the GA muscle of male Tak1^fl/fl^ and Tak1^mKO^ mice and differentially expressed genes (DEGs) in the RNA-seq dataset were identified using a threshold of Log2FC□≥□0.25 and *P* value□<□0.05. We found 2689 DEGs in GA muscle of Tak1^fl/fl^ and Tak1^mKO^ mice, of which 929 downregulated and 1760 upregulated (**Fig. 5A**). Pathway enrichment analysis using Metascape Annotation tool showed that upregulated genes in GA muscle of Tak1^mKO^ mice were associated with aerobic respiration and respiratory electron transport, mitochondrial gene expression, proteasome assembly, mitophagy, and positive regulation of protein catabolic processes (**Fig. 6B**) which is consistent with our previously published studies that genetic ablation of TAK1 augments proteolysis and accumulation of dysfunctional mitochondria in skeletal muscle (27).

**FIGURE 5.**
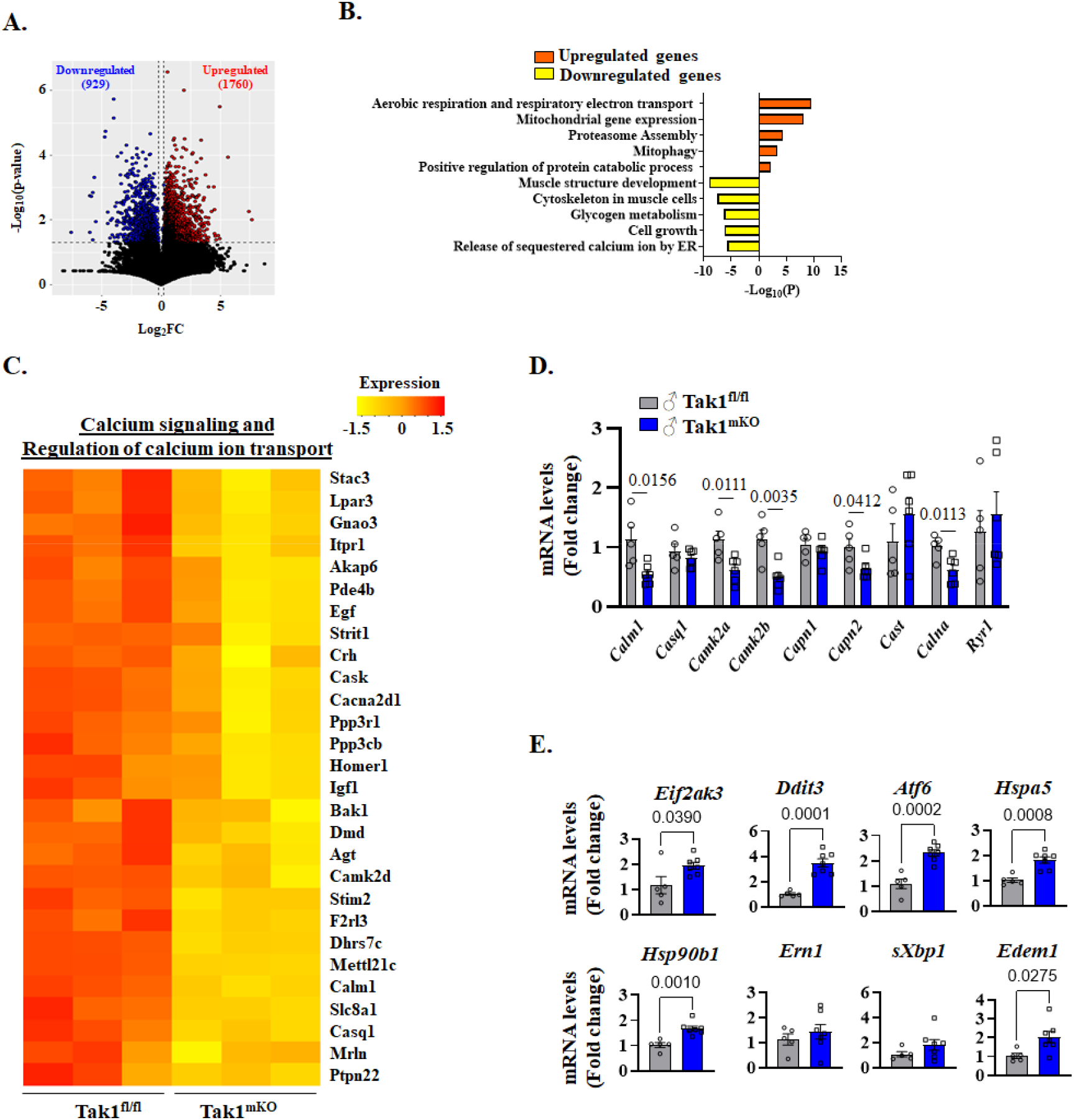
Role of TAK1 on the expression of genes related to calcium signaling and ER stress in skeletal muscle of male mice. **(A)**Volcano plot from RNA-Seq dataset analysis presented here shows differentially regulated genes (DEGs) in GA muscle of Tak1^fl/fl^ and Tak1^mKO^ male mice on day 30 after tamoxifen injection. n□=□3 mice per group. DEGs were identified with the threshold of Log2FC□≥□0.25 and *p*□<□0.05 using unpaired Student *t* test. **(B)** Bar graph shows pathways associated with downregulated and upregulated mRNAs in GA muscle of Tak1^mKO^ mice compared with Tak1^fl/fl^ mice identified using Metascape Analysis (metascape.org). **(C)** Heatmap showing relative mRNA levels of various molecules involved in calcium signaling and regulation of calcium ion transport in GA muscle of Tak1^fl/fl^ and Tak1^mKO^ mice. **(D)** Relative mRNA levels of calcium signaling associated molecules, *Calm1, Casq1, Camk2a, Camk2b, Capn1, Capn2, Cast, Calna*, and *Ryr1* in GA muscle of Tak1^fl/fl^ and Tak1^mKO^ male mice. **(E)** Relative mRNA levels of *Eif2ak3* (PERK), *Ddit3* (CHOP), *Atf6* (ATF6), Hspa5 (GRP78), Hsp90b1 (GRP94), *Ern1* (IRE1α), *sXbp1* (sXBP1), and *Edem1* (EBEM1) in GA muscle of Tak1^fl/fl^ and Tak1^mKO^ mice. n=5-7 mice per group. All data are presented as mean ± SEM. The data were analyzed with unpaired t-tests, and the resulting p-values are displayed in the graphs.

**FIGURE 6.**
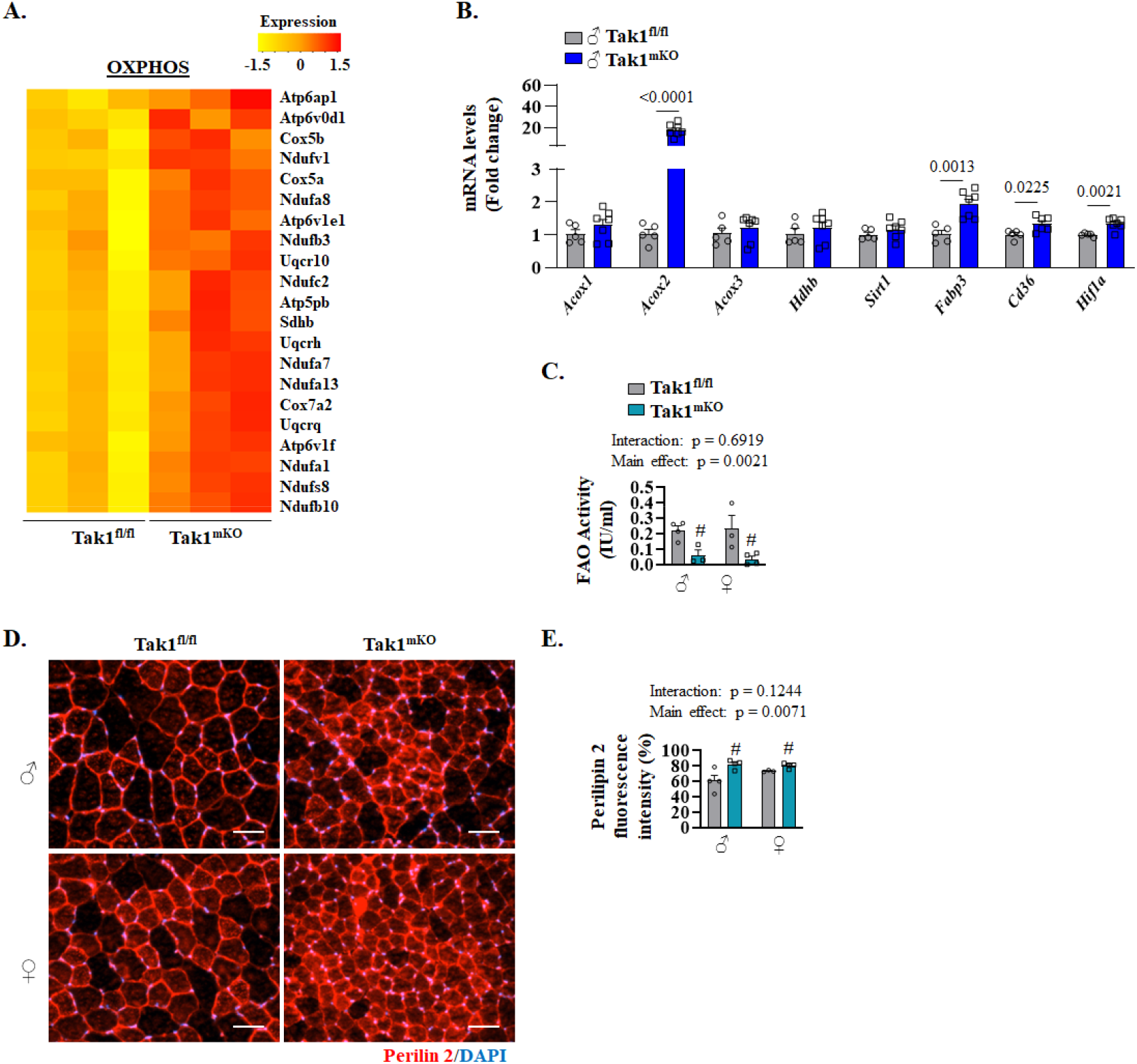
Role of TAK1 on fatty acid oxidation and lipid accumulation in skeletal muscle of male and female mice. **(A)**Heatmap showing relative mRNA levels of various components involved in oxidative phosphorylation (OXPHOS) in GA muscle of male Tak1^fl/fl^ and Tak1^mKO^ mice on day 30 after tamoxifen injection. **(B)** Relative mRNA levels of fatty acid metabolism-associated molecules (*Acox1, Acox2, Acox3, Hadhb, Sirt1, Fabp3, Cd36, Hif1a*) in GA muscle of male Tak1^mKO^ mice compared with male Tak1^fl/fl^ mice. n= 5-7 mice per group. Data are presented as mean ± SEM. The data were analyzed with unpaired t-tests, and the resulting p-values are displayed in the graphs. **(C)** Quantification of fatty acid oxidation (FAO) activity in GA muscle of male and female Tak1^fl/fl^ and Tak1^mKO^ mice. **(D)** Representative photomicrographs and **(E)** quantification of anti-Perilipin 2 fluorescence signal in TA muscle sections of male and female Tak1^fl/fl^ and Tak1^mKO^ mice. Scale bar, 50 μm. n=3-7 mice per group. All data are presented as mean ± SEM. The data were analyzed with two-way ANOVA and Tukey’s multiple comparison test, and the resulting p-values for the interactions, and main effect comparisions are displayed in the graphs.

Downregulated genes in GA muscle of Tak1^mKO^ were associated with muscle structure development, cytoskeleton in muscle cells, glycogen metabolism, cell growth, and release of sequestered calcium ions by ER (**Fig. 5B**). Heatmap generated from RNA-seq analysis further showed that gene expression of multiple molecules involved in calcium signaling were inhibited in Tak1^mKO^ mice compared with Tak1^fl/fl^ mice (**Fig. 5C**). Furthermore, independent qRT-PCR analysis showed that mRNA levels of various molecules, such as Calmodulin-1 (Calm1), Calcium/calmodulin-dependent protein kinase IIα and IIβ (Camk2a and Camk2b), Calpain-2 catalytic subunit (Capn2), and Calcineurin A-alpha (Calna), but not Calsequestrin-1 (Casq1), Calpastatin (Cast) and Ryanodine receptor 1 (Ryr1), were significantly reduced in GA muscle of male Tak1^mKO^ mice compared with Tak1^fl/fl^ mice (**Fig. 5D**). Since disruption in calcium signaling can lead to stress in endoplasmic reticulum ER (2, 30), we measured the expression of a few select molecules related to ER stress-induced UPR pathways. Interesting, there was a significant increase in the mRNA levels of Protein kinase R-like endoplasmic reticulum kinase (PERK, gene name: Eif2ak3), CHOP (Ddit3), ATF6, GRP78 (Hspa5), GRP94 (Hsp90b1), and Edem1 in GA muscle of male Tak1^mKO^ mice compared to corresponding male Tak1^fl/fl^ mice (**Fig. 5E**). These results suggest that muscle wasting in Tak1^mKO^ mice could be attributed to disruption in calcium homeostasis or signaling and associated ER stress.

### Targeted inactivation of TAK1 inhibits fatty acid oxidation in skeletal muscle of adult mice

Pathway analysis and heatmaps of DEGs in RNA-Seq dataset showed that gene expression of many molecules involved in aerobic respiration and oxidative phosphorylation or catabolic processes was increased in GA muscle of Tak1^mKO^ mice compared to Tak1^fl/fl^ mice (**Fig. 6A, Fig. S4**). Our independent qRT-PCR analysis further showed that mRNA levels of select molecules (e.g. Acox2, Fabp3, Cd36, and Hif1α) involved in fatty acid uptake or metabolism was significantly increased in GA muscle of male Tak1^mKO^ mice compared with male Tak1^fl/fl^ mice (**Fig. 6B**). Next, using a commercially available kit, we measured the fatty acid oxidation in GA muscle of male and female Tak1^fl/fl^ and Tak1^mKO^ mice. Interestingly, there was a significant reduction in the fatty acid oxidation in GA muscle of both male and female Tak1^mKO^ mice compared with corresponding Tak1^fl/fl^ mice (**Fig. 6C**). To understand whether TAK1 has any role in regulating the lipid droplet content in skeletal muscle, we performed immunostaining for the Perilipin 2 protein, which is one of the most abundantly expressed lipid droplet-coating proteins in skeletal muscle (31). Results showed that Perilipin 2 staining was significantly increased in TA muscle of both male and female Tak1^mKO^ mice compared to corresponding Tak1^fl/fl^ mice (**Fig. 6D, E**). These results suggest that TAK1 regulates fatty acid oxidation and lipid content in skeletal muscle of adult mice.

## Discussion

Our previous studies showed that TAK1 is essential for the maintenance of muscle mass in adult mice (27). However, it remained unknown whether there were any sex-related differences in the role of TAK1 in the regulation of muscle mass. In the present study, our results demonstrated that TAK1 is an important regulator of skeletal muscle mass, hypertrophic growth, and metabolic homeostasis in adult mice, with notable sex-dependent differences in the severity of muscle loss. Inactivation of TAK1 leads to rapid reductions in body weight and muscle mass, impaired overload-induced hypertrophy, altered lipid metabolism, and disruption of calcium signaling and ER homeostasis.

A key finding of this study is that TAK1 deletion induces more pronounced muscle atrophy in male mice than in female mice. While male Tak1^mKO^ mice exhibited significant reductions in body weight and in the mass of multiple hindlimb muscles, female Tak1^mKO^ mice showed a more restricted phenotype, with significant atrophy primarily in the TA muscle. Importantly, the relative reduction in myofiber CSA in the TA was similar in both sexes, suggesting that intrinsic myofiber atrophy occurs in males and females, but that female mice may possess compensatory mechanisms that partially preserve whole-muscle mass in certain muscle groups (**Fig. 1, 2**).

These sex differences may reflect the influence of sex hormones on anabolic and catabolic signaling pathways in skeletal muscle. Indeed, estrogen has been reported to exert protective effects against muscle wasting through modulation of inflammatory signaling, mitochondrial function, and oxidative stress (32-35). Therefore, it is possible that estrogen-dependent pathways mitigate the consequences of TAK1 loss in female mice. Alternatively, sex-specific differences in fiber-type composition, substrate utilization, or neuromuscular activity could also contribute to the differential sensitivity to TAK1 inactivation. Further studies examining hormonal manipulation or ovariectomy/orchidectomy models would help in understanding the mechanisms responsible for this sexual dimorphism.

Our results further demonstrated that TAK1 is activated in skeletal muscle in response to mechanical overload and it is required for overload-induced muscle growth in both male and female mice (**Fig. 3**). The absence of hypertrophy in Tak1-deficient plantaris muscle indicates that TAK1 functions as a central mediator of the signaling pathways that drive anabolic remodeling. Interestingly, the blunting of MOV-induced myofiber hypertrophy occurred without changes in the phosphorylation of Akt or mTOR but was associated with impaired phosphorylation of p70S6K and rpS6 protein. These findings suggest that TAK1 may regulate a branch of the translational machinery that converges on p70S6K and rpS6 independently of canonical Akt-mTOR signaling. These results are consistent with our previously published report which showed that forced activation of TAK1 increases protein synthesis in skeletal muscle of mice without increasing the phosphorylation of mTOR (28). Altogether, these findings suggest TAK1 functions in a parallel or complementary signaling pathway that is necessary for MOV-induced myofiber hypertrophy.

Another interesting finding of this study is the link between TAK1 and calcium-related signaling. RNA-seq and qRT-PCR analyses showed coordinated downregulation of multiple genes involved in calcium signaling, accompanied by increased expression of ER stress and UPR markers (**Fig. 6**). Because calcium signaling is essential for excitation-contraction coupling, mitochondrial function, and activation of anabolic pathways, its disruption is likely to have widespread consequences for muscle physiology. Chronic ER stress and activation of the UPR can promote muscle atrophy by inhibiting protein synthesis and enhancing proteolysis (2, 30, 36-38). Thus, the observed increase in ER stress markers in Tak1^mKO^ muscle provides a plausible mechanistic connection between TAK1 deficiency, impaired calcium handling, and muscle wasting.

Transcriptomic and biochemical analyses also revealed that TAK1 deletion profoundly alters metabolic gene expression and fatty acid utilization in skeletal muscle. Although genes associated with mitochondrial respiration and oxidative phosphorylation were upregulated, functional assays demonstrated reduced fatty acid oxidation and increased accumulation of lipid droplets, as indicated by enhanced Perilipin 2 staining (**Fig. 6)**. This apparent mismatch between gene expression and metabolic function suggests the presence of mitochondrial dysfunction or inefficient substrate utilization in Tak1-deficient muscle. Indeed, we have previously reported that oxygen consumption rate is significantly reduced in the isolated mitochondria from skeletal muscle of Tak1^mKO^ mice compared with Tak1^fl/fl^ mice further suggesting accumulation of dysfunctional mitochondria in Tak1-deficinet skeletal muscle (27, 29). Impaired fatty acid oxidation can lead to lipid accumulation, lipotoxic stress, and activation of proteolytic pathways, all of which contribute to muscle wasting (36, 39-41). The upregulation of genes involved in proteasome assembly and mitophagy further supports the notion that TAK1 loss triggers catabolic remodeling and turnover of damaged cellular components and that TAK1 plays an important role in maintaining metabolic flexibility and mitochondrial quality control in adult skeletal muscle.

Our study has several limitations that should be acknowledged. First, the RNA-seq analysis was performed exclusively on skeletal muscle from male Tak1^fl/fl^ and Tak1^mKO^ mice. Given the well-recognized sex-specific differences in skeletal muscle metabolism, signaling, and susceptibility to atrophy, inclusion of female mice in genome-wide expression and pathway analyses would provide a more comprehensive understanding of the role of TAK1 and help determine whether the observed transcriptional changes are sex-dependent. Second, although RNA-seq provides valuable insight into global transcriptional alterations, changes at the mRNA level do not necessarily reflect corresponding changes in protein abundance or activity. Therefore, additional validation of key differentially expressed genes using independent biochemical and molecular approaches and functional studies would further strengthen the mechanistic conclusions of this study.

In summary, our present study supports a model in which TAK1 integrates mechanical, metabolic, and stress-responsive signals to regulate skeletal muscle mass. Loss of TAK1 disrupts fatty acid oxidation, and calcium homeostasis while promoting ER stress and catabolic pathways, ultimately leading to myofiber atrophy and failure of hypertrophic growth. The partial protection observed in female mice highlights the importance of considering sex as a biological variable in studies of muscle signaling. Future studies will focus on defining the upstream activators and downstream effectors of TAK1 in skeletal muscle, as well as exploring its interactions with hormonal and metabolic regulators in both sexes.

## MATERIALS AND METHODS

### Animals

All mice were housed in a room maintained with a 12-12 h light-dark cycle and received food and water ad libitum. Wild-type C57BL6 male mice at 10 weeks of age were obtained from Jackson Laboratories. Tamoxifen inducible and skeletal muscle-specific Tak1 knockout mice were generated by crossing HSA-MCM mice (The Jackson Laboratory, Tg[ACTA1-cre/Esr1*]2Kesr/J) with Tak1^fl/fl^ mice as described (27, 28). All mice were in the C57BL/6 background, and their genotype was determined by PCR from tail DNA. For inactivation of TAK1, 10-12-week-old male and female mice were given i.p. injections of tamoxifen (75 mg/kg body weight) in corn oil for 4 consecutive days, and the mice were kept on tamoxifen-containing standard chow (Harlan Laboratories) (250 mg/kg) for the entire duration of the experiment. Body weights were recorded on day 15 and 30 after tamoxifen injections.

### Bilateral mechanical overload (MOV) surgery

10-week-old male and female Tak1^fl/fl^ and Tak1^mKO^ were treated with tamoxifen for 4 consecutive days and after a 2-day washout period, mice underwent sham or bilateral MOV surgery. In brief, adult Tak1^fl/fl^ and Tak1^mKO^ male and female mice were anesthetized using isoflurane, followed by making a ∼0.5-cm incision proximal to the ankle, separating the tendons, and removing the solues along with the distal tendon and associated myotendious junction of the GA while keeping the plantaris muscle intact in both limbs. A sham surgery was performed for control mice following the same procedures, except that tendons of the GA and soleus muscles were kept intact. The incisions were closed with surgical sutures. Finally, the mice were euthanized on day 7 or 14 after MOV surgery, and the plantaris muscles were collected for biochemical and histological analyses. All animal procedures were conducted in strict accordance with the institutional guidelines and were approved by the Institutional Animal Care and Use Committee and Institutional Biosafety Committee of the University of Houston (PROTO201900043).

### Histology, immunohistochemistry, and morphometric analysis

Individual TA, soleus, or plantaris muscle was isolated from mice, snap-frozen in liquid nitrogen, and sectioned with a microtome cryostat. For the assessment of muscle morphology, 8-μm-thick transverse sections of TA and plantaris muscle were stained with hematoxylin and eosin (H&E) dye and examined under Nikon Eclipse Ti-2E Inverted Microscope (Nikon). Muscle sections were also processed for immunostaining for dystrophin protein to mark the boundaries of myofibers. For immunostained sections, the slides were mounted using fluorescence medium (Vector Laboratories) and visualized at room temperature on Nikon Eclipse Ti-2E Inverted Microscope (Nikon), a digital camera (Digital Sight DS-Fi3, Nikon), and Nikon NIS Elements AR software (Nikon). For morphometric analysis, myofibers were selected randomly across the entire muscle cross-section to avoid regional sampling bias. Average myofiber cross-sectional area (CSA) was determined in anti-dystrophin-stained muscle sections using ImageJ software (NIH, Bethesda, MD). Approximatly 200 myofibers per muscle per mouse were analysed. All image acquisition and CSA measurements were performed in a blinded manner. A frequency distribution curve was generated using categorical intervals of 400 μm^2^ myofiber CSA.

### Immunostaining for Perilipin 2 protein

To assess the abundance of intramuscular lipid droplets, Perilipin 2 immunostaining was performed on TA muscle transverse sections as described (36). Briefly, the muscle transverse cryosections were fixed in 4% PFA in PBS for 15□min, followed by rinsing with PBS and blocking with 3% BSA solution. Subsequently, muscle sections were incubated with Perilipin 2 (Plin2) antibody (Proteintech, Rosemont, IL) overnight at 4□°C in a humidified chamber. The Perilipin 2 staining was detected using AF555 anti-rabbit IgG. Nuclei were counterstained with DAPI. The Plin2 fluorescence intensity was quantified using ImageJ software (NIH, Bethesda, MD).

### Measurement of fatty acid oxidation (FAO)

FAO activity in skeletal muscle was measured using the fatty acid oxidation (FAO) assay kit (Biomedical Research Service, SUNY Buffalo, US) following the manufacturer’s suggested protocol.

### RNA extraction and qRT-PCR

RNA isolation and qRT-PCR were performed following a protocol as described (36, 42). In brief, total RNA was extracted from the GA muscle of Tak1^fl/fl^ and Tak1^mKO^ mice using TRIzol reagent (ThermoFisher Scientific) and RNeasy Mini Kit (Qiagen, Valencia, CA, USA) according to the manufacturer’s protocols. First-strand cDNA for PCR analysis was made with a commercially available kit (iScript cDNA Synthesis Kit, Bio-Rad Laboratories). The quantification of mRNA expression was performed using the SYBR Green dye (Bio-Rad SsoAdvanced-Universal SYBR Green Supermix) method on a sequence detection system (CFX384 Touch Real-Time PCR Detection System, Bio-Rad Laboratories). The sequence of the primers is described in Supplemental **Table S1**. Data normalization was accomplished with the endogenous control (GAPDH), and the normalized values were subjected to a 2-ΔΔCt formula to calculate the fold change between control and experimental groups.

### Western blot

GA or plantaris muscle of mice was homogenized in lysis buffer (50□mM Tris-Cl (pH 8.0), 200□mM NaCl, 50□mM NaF, 1□mM dithiothreitol, 1□mM sodium orthovanadate, 0.3% IGEPAL, and protease inhibitors). Approximately 100□μg protein of whole protein lysate was resolved on each lane on 8-12% SDS-PAGE gel, transferred onto a nitrocellulose membrane, and probed using a specific primary antibody. Bound antibodies were detected by secondary antibodies conjugated to horseradish peroxidase (Cell Signaling Technology). Signal detection was performed by an enhanced chemiluminescence detection reagent (Bio-Rad). Approximate molecular masses were determined by comparison with the migration of prestained protein standards (Bio-Rad). Antibodies and their dilution ratios used in the study are provided in Supplemental **Table S2**. The uncropped gel images for western blot are presented in Supplemental **Fig. S5**.

### RNA-seq and data analysis

RNA-seq experiment and data analysis were performed following a method as described (42). In brief, the mRNA-seq library was prepared using poly (A)-tailed enriched mRNA at the UT Cancer Genomics Center using the KAPA mRNA HyperPrep Kit protocol (KK8581, Roche, Holding AG, Switzerland) and KAPA Unique Dual-indexed Adapter kit (KK8727, Roche). The Illumina NextSeq550 was used to produce 75 base paired-end mRNA-seq data at an average read depth of ∼38□M reads/sample. RNA-seq fastq data were processed using CLC Genomics Workbench 20 (Qiagen). Illumina sequencing adapters were trimmed, and reads were aligned to the mouse reference genome Refseq GRCm39.105 from the Biomedical Genomics Analysis Plugin 20.0.1 (Qiagen). Normalization of RNA-seq data was performed using a trimmed mean of M values. Genes with Log2FC□≥□□0.25□ and P value□<□0.05 were assigned as differentially expressed genes (DEGs) and represented in a volcano plot using ggplot function in R software (v 4.2.2). Pathway enrichment analysis was performed using the Metascape Gene Annotation and Analysis tool (metascape.org) as described (Zhou et al, 2019). Heatmaps were generated by using heatmap.2 function (Gu and Hübschmann, 2022) using z-scores calculated based on transcripts per million (TPM) values. TPM values were converted to log (TPM□□+□□1) to handle zero values. Genes involved in specific pathways were manually selected for heatmap expression plots.

### Statistical Analysis

All wet-lab data are presented as mean ± standard error of the mean (SEM). Statistical analyses were performed using GraphPad Prism 10.0. Comparisons between two groups were conducted using a two-tailed unpaired Student’s t-test. For experiments involving multiple comparisons, two-way analysis of variance (ANOVA) followed by Tukey’s post hoc test was applied. *p* ≤ 0.05 was considered statistically significant. Additional statistical details, including exact n, are provided in the corresponding figure legends.

## Authors’ contributions

AK and TAH designed the work. MTS, ASJ, and AR performed the experiments and analyzed the results. MTS and ASJ wrote the first draft of the manuscript. AK, TAH and other authors edited and finalized the manuscript.

### Acknowledgements

We are highly grateful to Dr. S. Akira (Osaka University, Osaka, Japan) for providing floxed TAK1 mice.

