## Supplementary material for "TAK1 regulates skeletal muscle mass, hypertrophic signaling, and metabolic homeostasis in male and female mice": Figures S1-5, Tables S1 and S2

This file contains **Figures S1-S5** and **Tables S1 and S2**

### Supplementary Figures and Legends

**FIGURE S1**

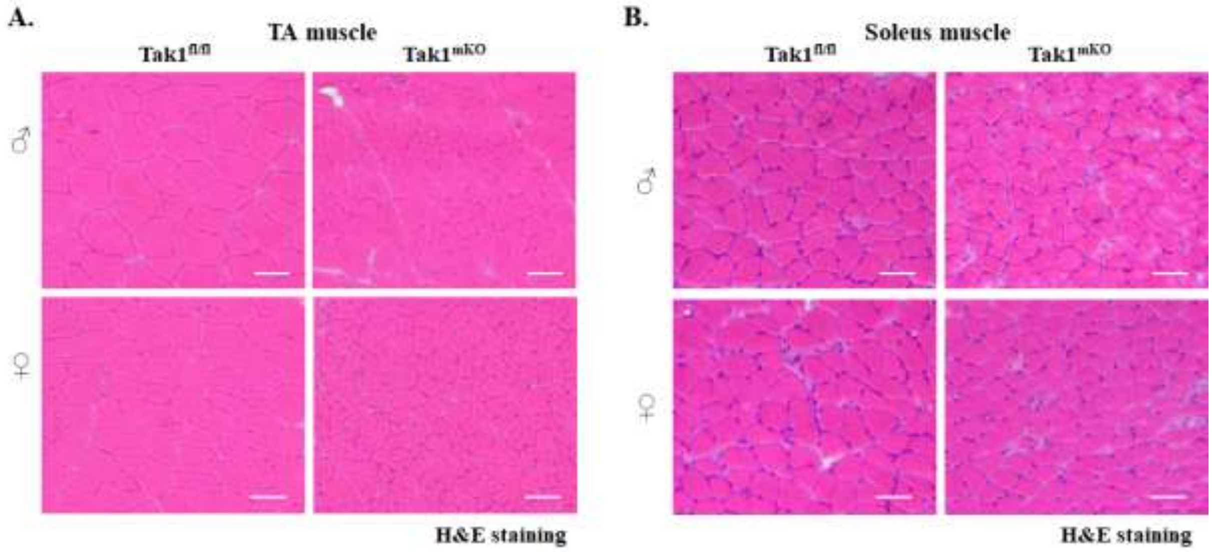


**Figure S1. Effect of TAK1 inactivation of muscle structure in male and female Tak1$^{fl/fl}$ and Tak1$^{mKO}$ mice.** Representative Hematoxylin and Eosin (H&E)-stained images of **(A)** tibialis anterior (TA), and **(B)** soleus (SOL) muscle of male and female Tak1$^{fl/fl}$ and Tak1$^{mKO}$ mice after 30 days of start of tamoxifen injections. Scale bar, 50 µm.

FIGURE S2

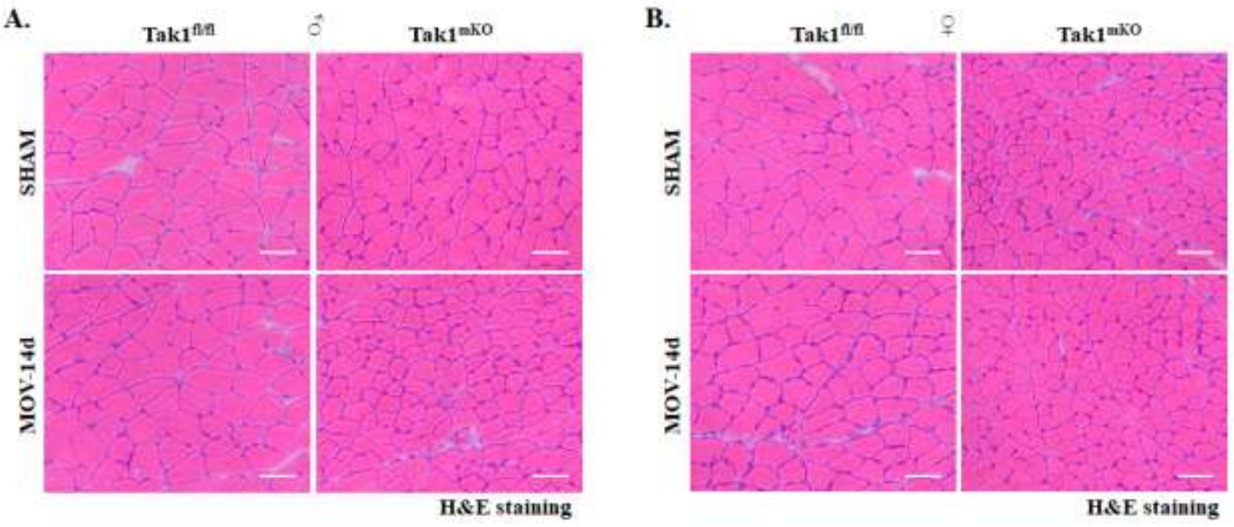


**Figure S2. Effect of TAK1 inactivation of muscle structure of plantaris muscle of male and female Tak1$^{fl/fl}$ and Tak1$^{mKO}$ mice.** Representative Hematoxylin and Eosin (H&E)-stained images of plantaris muscle of **(A)** male, and **(B)** female Tak1$^{fl/fl}$ and Tak1$^{mKO}$ mice after 14 days of performing sham or bilateral MOV surgery. Scale bar, 50 µm.

**FIGURE S3**

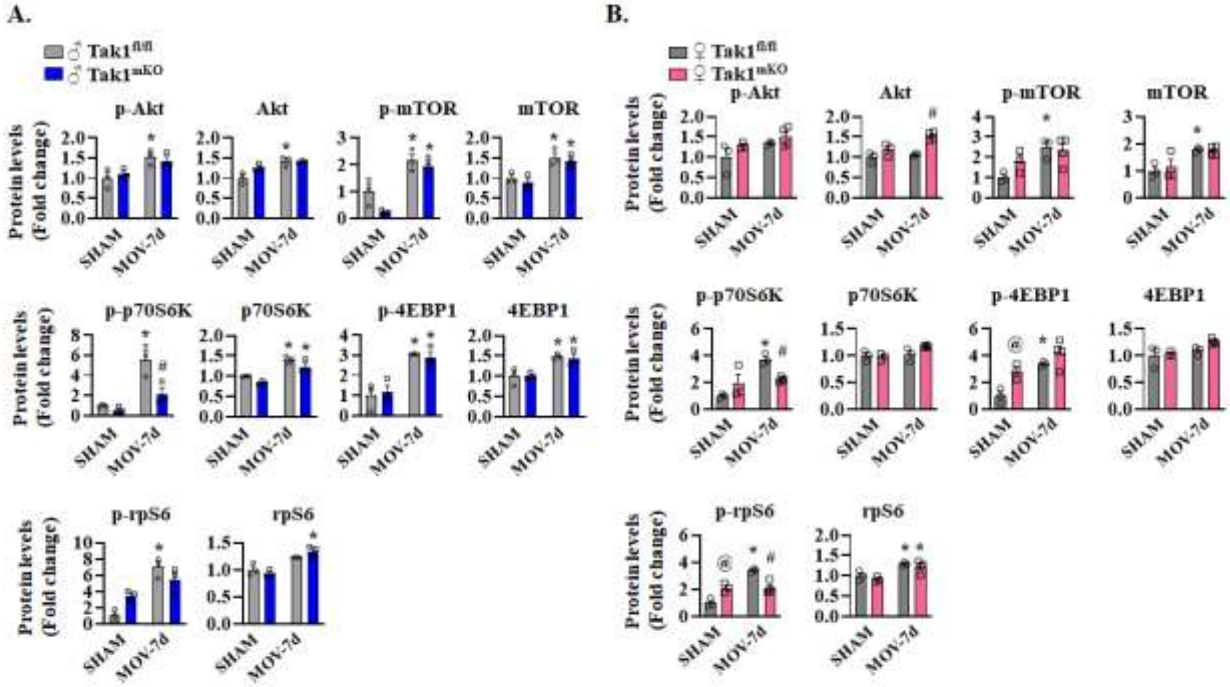


**Figure S3. Effect of targeted inactivation of TAK1 on the phosphorylation of the components of Akt-mTOR signaling in male and female mice.** Densitometry analysis of phosphorylated and total levels of Akt, mTOR, p70S6K, 4EBP1, and Rps6 proteins in plantaris muscle of **(A)** male **(B)** female Tak1$^{fl/fl}$ and Tak1$^{mKO}$ mice on day 7 after performing sham or MOV surgery. n=3-4 mice per group. All data are presented as mean ± SEM. $^{*}p < 0.05$, values significantly different from corresponding sham operated mice; $^{\#}p < 0.05$, values significantly different from Tak1$^{fl/fl}$ mice subjected to MOV surgery; $^{@}p < 0.05$, values significantly different from female Tak1$^{fl/fl}$ sham operated mice analyzed by two-way ANOVA and Tukey's multiple comparison test.

### FIGURE S4

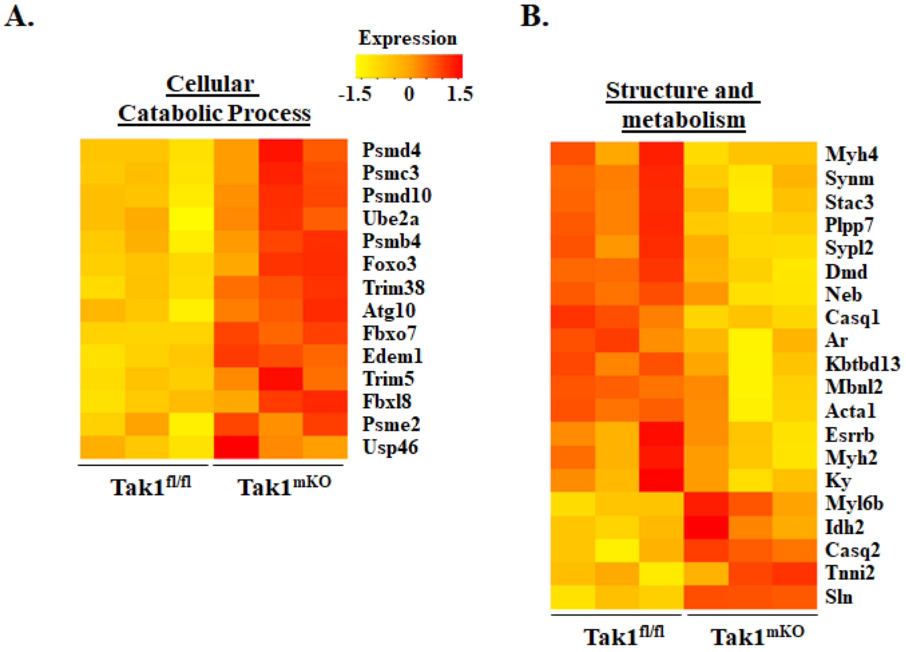


**Figure S4. Effect of TAK1 inactivation on the gene expression of molecules associated with catabolic processes and muscle structural integrity.** Heatmap showing relative mRNA levels of multiple molecules associated in **(A)** cellular catabolic process and **(B)** muscle structure and metabolism in GA muscle of $Tak1^{fl/fl}$ and $Tak1^{mKO}$ mice.

**FIGURE S5**

**1F.**

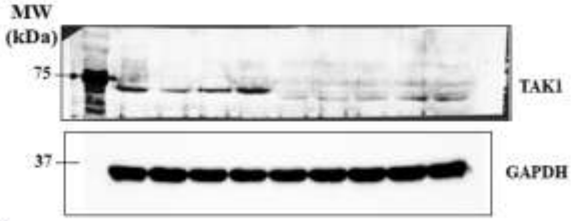


**1G.**

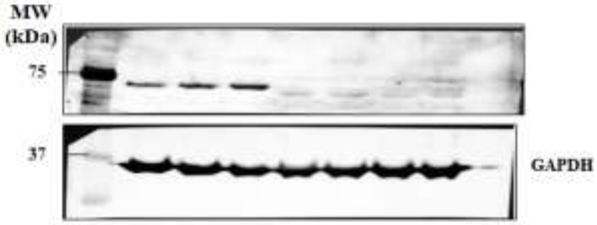


**3A.**

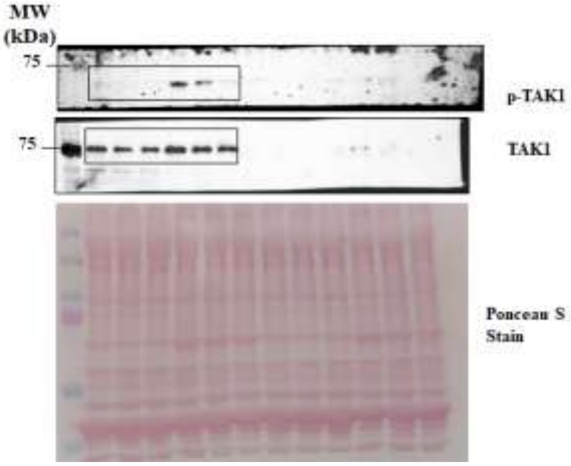


**3C.**

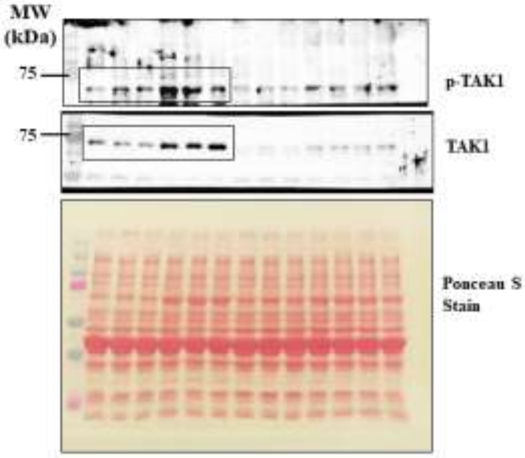

FIGURE S5 (cont.)

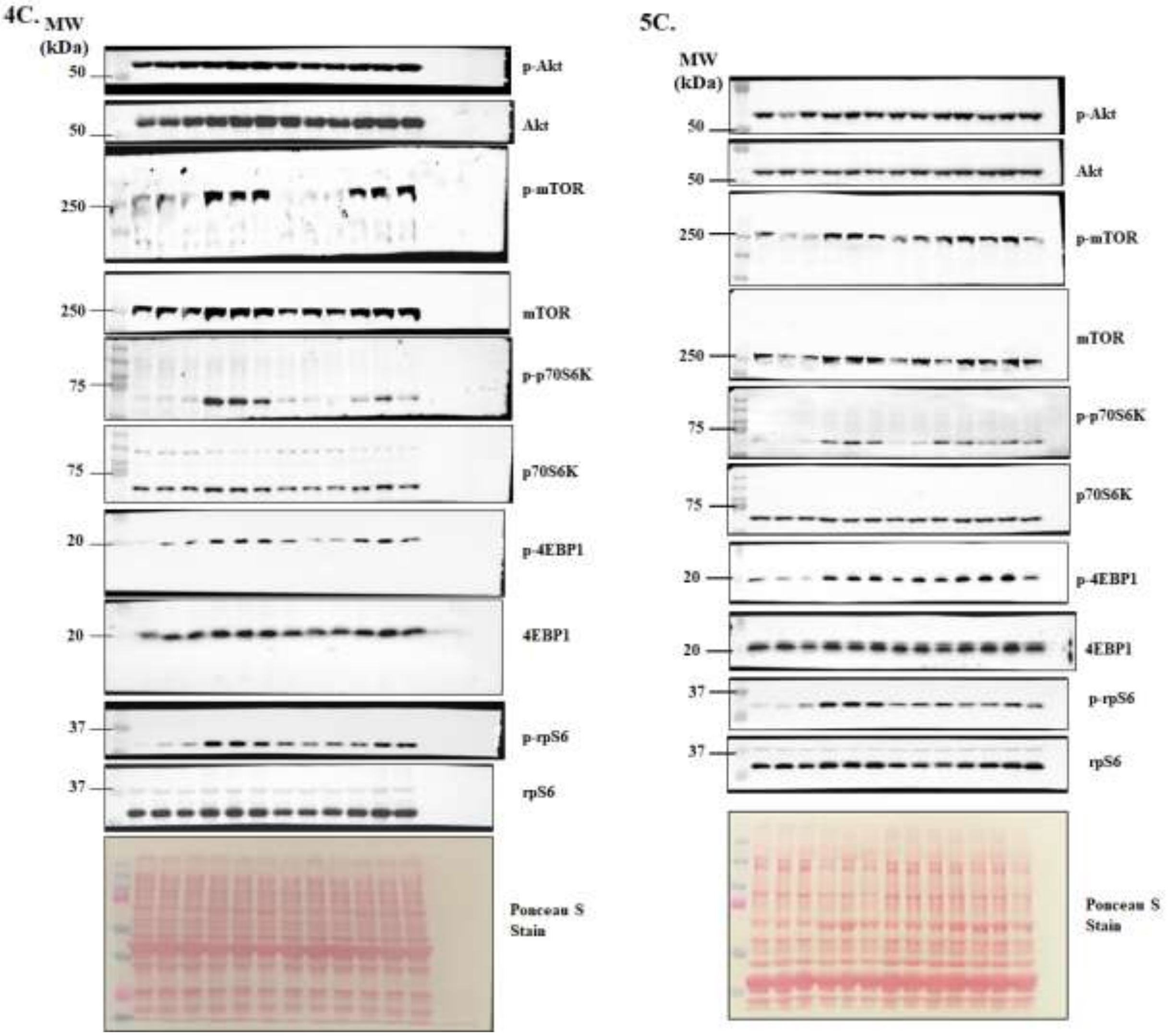


**Figure S5. Uncropped gel images.** Uncropped immunoblots used for various experiments in the main manuscript file.

**Table S1. Sequence of the primers used in qRT-PCR assay.**

| Gene Name | Forward primer (5'-3') | Reverse primer (5'-3') |
|---|---|---|
| Acox1 | GGATGGTAGTCCGGAGAACA | AGTCTGGATCGTTCAGAATCAAG |
| Acox2 | CCTTCCTAGACCTGCTTCCC | TGTCCGTCATAACAGCCAAG |
| Acox3 | CTTCTGAGAAACGGGGACAA | GCTCGGTAGGCACTAAGAGG |
| Hadhb | GCACTTTCGGGTTTGTTG | GTGTGAGCTGGAGTCTTATC |
| Sirt1 | GACGATGACAGAACGTCACAC | CGAGGATCGGTGCCAATCA |
| *Fabp3* | CCCCTCAGCTCAGCACCAT | CAGAAAAATCCCAACCCAAGAAT |
| Cd36 | GGCCAAGCTATTGCGACAT | CAGATCCGAACACAGCGTAGA |
| Hif1a | TGAGCTTGCTCATCAGTTGC | CCATCTGTGCCTTCATCTCA |
| Calm1 | TGGGAATGGTTACATCAGTGC | CGCCATCAATATCTGCTTCTCT |
| *Casq1* | CTGGCACTGCTGTTTGTACTG | GGGGGCTCATGGTAGAGGAG |
| *Camk2a* | TGGAGACTTTGAGTCCTACACG | CCGGGACCACAGGTTTTCA |
| *Camk2b* | CGTTTCACCGACGAGTACCAG | GCGTACAATGTTGGAATGCTTC |
| *Capn1* | ATGACAGAGGAGTTAATCACCCC | GCCCGAAGCGTTTCATAATCC |
| *Capn2* | GGAGAGAGGCTGTACCTTCCT | CCGAGGTGGATGTTGGTCTG |
| *Cast* | GGAAGGACAAACCAGAGAAGC | AGGGGCAGCTATCCAAATCTT |
| *Calna* | GTGAAAGCCGTTCCATTTCCA | GAATCGAAGCACCCTCTGTTATT |
| *Ryr1* | CAGTTTTTGCGGACGGATGAT | CACCGGCCTCCACAGTATTG |
| *Eif2ak3* | ACT CCT GTC TTG GTT GGG TCT GAT | CGT GCT CCG CTT ATT CCT TTC T |
| *Ddit3* | TGAAAGCAGAACCTGGTCCA | CACTGTTCATGCTTGGTGCA |
| *Atf6* | CGTTCCTGAGGAGTTGGATTTG | GCTTCTCTTCCTTCAGTGGCTCTA |
| *Hspa5* | CGT GGA GAT CAT AGC CAA CGA T | ATT CCA AGT GCG TCC GAT GA |
| *Hsp90b1* | GGG AGG TCA CCT TCA AGT CG | CTC GAG GTG CAG ATG TGG G |
| *Ern1* | CCTTTGCTGATAGTCTCTGCCCAT | TTACCACCAGTCCATCGCCATT |
| *sXbp1* | AAGAACACGCTTGGGAATGG | CTGCACCTGCTGCGGAC |
| *Edem1* | CGGCTATGACAACTACATGG | GTTCAGATTGGACTCTC |
| β-actin | CAGGCATTGCTGACAGGATG | TGCTGATCCACATCTGCTGG |

**Table S2. List of the antibodies used for immunofluorescence (IF) staining or western blot (WB).**

| Antibody | Source and Catalog no. | Dilution IF/WB |
|---|---|---|
| Monoclonal rabbit-anti-total-TAK1 | Cell Signaling Technology, # 5206 | 1:500 |
| Monoclonal rabbit-anti-GAPDH | Cell Signaling Technology # 2118 | 1:500 |
| Polyclonal rabbit-anti-Dystrophin | Abcam, ab15277 | 1:250 |
| Donkey anti-Rabbit IgG Alexa Fluor 555 | Invitrogen # A31572 | 1:1000 |
| Monoclonal rabbit-anti-phospho-Akt (Ser473) | Cell Signaling Technology # 4060 | 1:1000 |
| Polyclonal rabbit-anti-total-Akt | Cell Signaling Technology # 9517 | 1:1000 |
| Polyclonal rabbit-anti-phospho-mTOR (Ser2448) | Cell Signaling Technology # 2971 | 1:500 |
| Polyclonal rabbit-anti-total-mTOR | Cell Signaling Technology # 2972 | 1:500 |
| Polyclonal rabbit-anti-phospho-p70S6K (Thr389) | Cell Signaling Technology # 9205 | 1:500 |
| Polyclonal rabbit-anti-p70S6K | Cell Signaling Technology # 9202 | 1:500 |
| Monoclonal rabbit-anti-phospho-4E-BP1 (Thr37/46) | Cell Signaling Technology # 2855 | 1:500 |
| 4E-BP1 (53H11) Rabbit | Cell Signaling Technology # 9644 | 1:500 |
| Monoclonal rabbit-anti-phospho-rpS6 (Ser235/236) | Cell Signaling Technology # 4858 | 1:500 |
| Monoclonal rabbit-anti-rpS6 | Cell Signaling Technology # 2217 | 1:500 |
| Monoclonal rabbit-anti-phospho-TAK1 (Thr184, Thr187) | Invitrogen #MA5-15073 | 1:500 |
| Rabbit anti-Perilipin-2 | Proteintech # 15294-1-AP | 1:300 |
| Anti-rabbit IgG HRP | Cell Signaling # 7074S | 1:2000 |
| Anti-mouse IgG HRP | Cell Signaling # 7076S | 1:2000 |
